# Nicotine stimulates chemosensory behavior and reduces aggression towards tobacco-feeding caterpillars (*Manduca sexta*) by predatory side-blotched lizards (*Uta stansburiana*)

**DOI:** 10.64898/2026.11.24.768842

**Authors:** Caroline A. Pete, Sarah E. Pottinger, Ian T. Baldwin, Jay K. Goldberg

**Author notes:** **Author for correspondence:** Jay K Goldberg.

## Abstract

Herbivorous insects that feed on toxic plants have not only evolved to tolerate the chemical defenses of their hosts, but to co-opt them for their own defenses. Previous research has shown that the tobacco hornworm (*Manduca sexta*), a specialist herbivore of alkaloid-rich Solanaceous plants, is capable of exhaling ingested nicotine to defend itself from spiders; a phenomenon known as defensive halitosis. In this study, we aimed to uncover the effect of nicotine on interactions between predatory lizards (*Uta stansburiana*) and lepidopteran larvae (*Manduca sexta* and *Spodoptera exigua*). We leveraged transgenic lines of wild tobacco (*Nicotiana attenuata*) that are unable to produce nicotine or that inhibit *M. sexta*’s defensive halitosis via RNAi. We found that ingested nicotine in prey altered the rates of chemosensory and aggressive behaviors performed by lizards, yet there was no effect of inhibiting *M. sexta*’s ability to use ingested nicotine in a defensive halitosis. Furthermore, we found that nicotine had no effect on interactions between lizards and *S. exigua* larvae. These results suggest that nicotine on the surface of tobacco-feeding insects, but not volatile nicotine, influences lizard behavior in a dose-dependent manner rather than being a universally aversive chemical cue.

## Introduction

Many herbivorous insects have evolved to not only tolerate chemically defended host plants, but to co-opt plant defenses for their own benefit. Sequestration of plant chemistry is a widespread phenomenon and has been documented in numerous insect species that feed on a diverse array of host plants (Beran and Petschenka 2022). Sequestered plant compounds can defend herbivores through a variety of mechanisms including direct toxicity (Narberhaus et al 2005) and olfactory camouflage (Bui et al 2021). The plant chemicals that are co-opted by herbivores are as diverse as the insects that use them and can protect against an equally diverse range of natural enemies, including parasitoids and insectivores (Petschenka and Agrawal 2016).

Lizards are major insectivores and often rely heavily on their chemical senses for numerous life processes, especially when foraging for arthropod prey (Cooper 1997). For some especially olfactory species, chemical cues of prey presented on cotton swaps are enough to stimulate aggressive feeding behaviors (e.g. biting swabs; Cooper 1995a; Goldberg et al. 2021). The readily observed chemosensory (e.g. tongue flicks) and aggressive behaviors of lizards facilitates the study of their responses to various chemical stimuli (Cooper 1994), including isolated plant-derived chemicals which have been shown to elicit behavioral responses from predatory lizards (Stork et al. 2011; Goldberg et al. 2019).

Nicotine, well known for its psychoactive properties in humans (Pomerleau 1992), functions as a chemical defense in wild plants (Steppuhn et al 2004). It is primarily produced by members of the genus *Nicotiana* (Solanales: Solanaceae; Xu et al 2017) and has a strong affinity for the acetylcholine receptors of animals (Santis et al. 2024). This affinity is the primary mechanism for nicotine’s broad toxicity, which enables it to defend plants against a wide range of herbivores (Steppuhn et al 2004). It is biosynthesized via the conjugation of nicotinic acid and N-methylpyrrolinium (Chang et al. 2026). A key step in the synthesis of N-methylpyrrolinium is carried out by putrescine methyl-transferase (PMT), which converts putrescine into N-methylputrescine (Chang et al. 2026). The loss of PMT function results in nicotine-free plants, thereby enabling studies of nicotine’s defensive role while keeping biological context (ie: plant physiology) otherwise intact (Steppuhn et al 2004).

In humans, nicotine is metabolized by enzymes in the liver to a variety of compounds, predominantly cotinine (Benowitz et al. 2009). Some generalist insect herbivores metabolize nicotine through a similar pathway, while tobacco-specializing herbivores have evolved other strategies to achieve nicotine tolerance (Kumar et al 2014a). *Manduca sexta*, for example, has been shown to produce no nicotine oxides after feeding on nicotine-rich plants (Kumar et al 2014a/b). Furthermore, *M. sexta* is able to exhale volatilized nicotine from its spiracles to deter predatory spiders (Kumar et al 2014a/b). This “defensive halitosis” is mediated by nicotine-induced expression of the gene *MsCYP6B46* (hereafter CYP) in larval midguts; however, it does not deter all predators that *M. sexta* larvae encounter in nature (Kumar et al. 2014a/b).

We set out to examine the responses of predatory side-blotched lizards (*Uta stansburiana*) to lepidopteran larvae that had been fed on nicotine-free and nicotine-containing plants. By leveraging multiple stable transgenic lines of wild tobacco (*Nicotiana attenuata*) and readily accessible lepidopteran species (*Manduca sexta* and *Spodoptera exigua*), we addressed three overarching questions: 1) how does nicotine impact lizard feeding behavior? 2) How do lizards respond to herbivore species with different nicotine-metabolism pathways? and 3) Does CYP-mediated nicotine exhalation impact interactions between lizards and *M. sexta* larvae? We found that all three factors – nicotine, prey species, and CYP-mediated nicotine exhalation – impacted lizard responses to prey, especially rates of chemosensory and aggressive (e.g. biting) behaviors.

## Methods

### Study site and species

Trials were conducted at the Lytle Ranch Preserve in Santa Clara, Utah, 84765 (37°08′45′′N, 114°01′11′′W), well within the native ranges of both *Nicotiana attenuata* and *Manduca sexta*. The common side-blotched lizard (*Uta stansburiana*) – a generalist predator of small herbivores that readily feeds on lepidopteran larvae (Figure 1) – is common in this area. Stable transgenic lines of *N. attenuata* were generated via *Agrobacterium tumefaciens*-mediated transformation as previously described (Krugel et al. 2002). Nicotine-free irPMT plants were generated by using RNAi to reduce PMT expression and prevent nicotine biosynthesis (Steppuhn et al. 2004); irCYP plants were used to silence the expression of MsCYP6B46 in *M. sexta* larvae (Kumar et al. 2012); and an empty vector (EV) control was also used (Schwachtje et al. 2008). Transgenic *N. attenuata* seeds were imported and released in accordance with several Animal and Plant Health Inspection Service notifications (detailed list previously published in Kumar et al. 2014).

**Figure 1.**
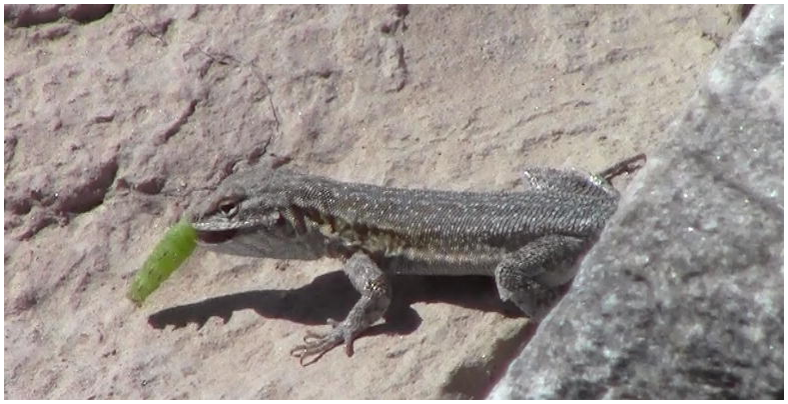
Still image of a lizard (*Uta stansburiana*) consuming an EV-fed *Manduca sexta* larvae during a field behavior trial.

### Caterpillar rearing

All *N. attenuata* plants were grown outdoors on the Lytle Ranch Preserve as previously described (Kessler et al. 2012). Freshly hatched *Manduca sexta* neonates were assigned to a treatment (EV, irCYP, or irPMT) and placed on the corresponding *N. attenuata* leaves in a small plastic cup. Freshly picked leaves were provided daily, while old leaves were removed. *Manduca sexta* larvae were allowed to feed for 14 days before being used in lizard trials. *Spodoptera exigua* larvae fed on artificial diet (Kumar et al 2014a) for 14 days, before being switched to *N. attenuata* leaves on which they fed for four additional days before being used in lizard trials. Detailed rearing conditions are described in (Kumar et al 2014b). The irPMT + nicotine caterpillars were sprayed with a 1mg/mL solution of analytical-grade nicotine in distilled water and allowed to dry on paper towels immediately before trials began.

### Behavioral Trials

All trials (N = 45) occurred between 0900 and 1200 from May 31 to June 4, 2014. Lizards basking on rocks or logs in the Beaver Dam Wash were identified by an individual observer (JKG) who recorded all trials from a distance of ∼3m. Prey caterpillars were tossed 30-50cm in front of the lizard. Trials began if/when a lizard physically interacted with the prey (usually by nose-tapping, tongue-flicking, or biting it) and continued until 30 seconds after a lizard had completely ingested the prey or had lost interest (defined as moving >50cm away for >1min). Lizards were never captured, handled, or interacted with beyond the offering of prey during experimental trials.

### Ethogram and behavior scoring

An observer with no prior knowledge of the treatment conditions (SEP) quantified five lizard behaviors from video recordings (published in Goldberg and Baldwin 2026): tongue-flicks, a chemosensory behavior in which animals briefly tap their tongue on an object to pass odorants to their olfactory and vomeronasal systems (Cooper 1994); lip licks, a chemosensory behavior similar to tongue-flicks but directed towards the air (Goldberg et al 2019); head bobs, a display behavior directed at conspecifics and potential predators (Martins et al 2004); bites directed at the prey; and the duration of thrashing bouts in which the lizard thrashes wildly while prey is in its jaws. The rate at which each behavior was observed (occurrences per second) was then calculated in order to account for the variation in trial length resulting from our design (min = 38 seconds; max = 1687 seconds; mean = 262.8 seconds). For thrashing behavior, the total duration of all bouts was added together and standardized for trial length by calculating the proportion of time performing the behavior (time spent thrashing/total trial duration). The observer also noted the final outcome of the trial (prey eaten/uneaten).

### Statistical analyses

All statistical analyses for this investigation were performed using the R statistical computing software (R Core Team 2025). Each of our two prey species (*Manduca sexta* and *Spodoptera exigua*) were treated as separate experiments and analyzed independently. We used a binomial generalized linear modeling approach to analyze differences between treatments in the proportion of trials resulting in eaten or uneaten prey. Other behavioral variables were analyzed using a one-way ANOVA or Kruskal-Wallis test, depending on whether or not they satisfied the assumptions of parametric tests. For variables with significant omnibus tests, pairwise *post-hoc* analyses were then employed (Tukey’s HSD for ANOVA; Conover Test for Kruskal-Wallis). A significance cutoff of p=0.05 was used in all analyses.

## Results

### Nicotine’s effect on caterpillar consumption

We found no significant effects of nicotine on the likelihood of lizards to consume their prey (logit glm; all coefficients found to have p>0.05); however, there are clear differences in the likelihood of lizards to consume prey of different treatment types, indicating that our lack of significance is at least in part due to the low statistical power of the small sample size. No irPMT+N *M. sexta* caterpillars were eaten by lizards, whereas varying proportions of other treatments were eaten (55.6% of EV, 33.3% of irCYP, 75% irPMT; Table 1; Figure 2). On the other hand, all *Spodoptera exigua* larvae were consumed during behavior trials (Table 1). Given that *S. exigua* larvae are significantly smaller than *M. sexta* larvae, the difference in lizard consumption rates between prey species may be driven by caterpillar size.

**Table 1.** Proportion of prey that were eaten during trials.

| PREY SPECIES | TREATMENT | TOTAL TRIALS | PREY EATEN | %EATEN |
|---|---|---|---|---|
| *M. SEXTA* | EV | 9 | 5 | 55.6% |
| *M. SEXTA* | CYP | 12 | 4 | 33.3% |
| *M. SEXTA* | PMT | 8 | 6 | 75% |
| *M. SEXTA* | PMT+N | 6 | 0 | 0% |
| *S. EXIGUA* | EV | 2 | 2 | 100% |
| *S. EXIGUA* | PMT | 4 | 4 | 100% |
| *S. EXIGUA* | PMT+N | 5 | 5 | 100% |

**Figure 2.**
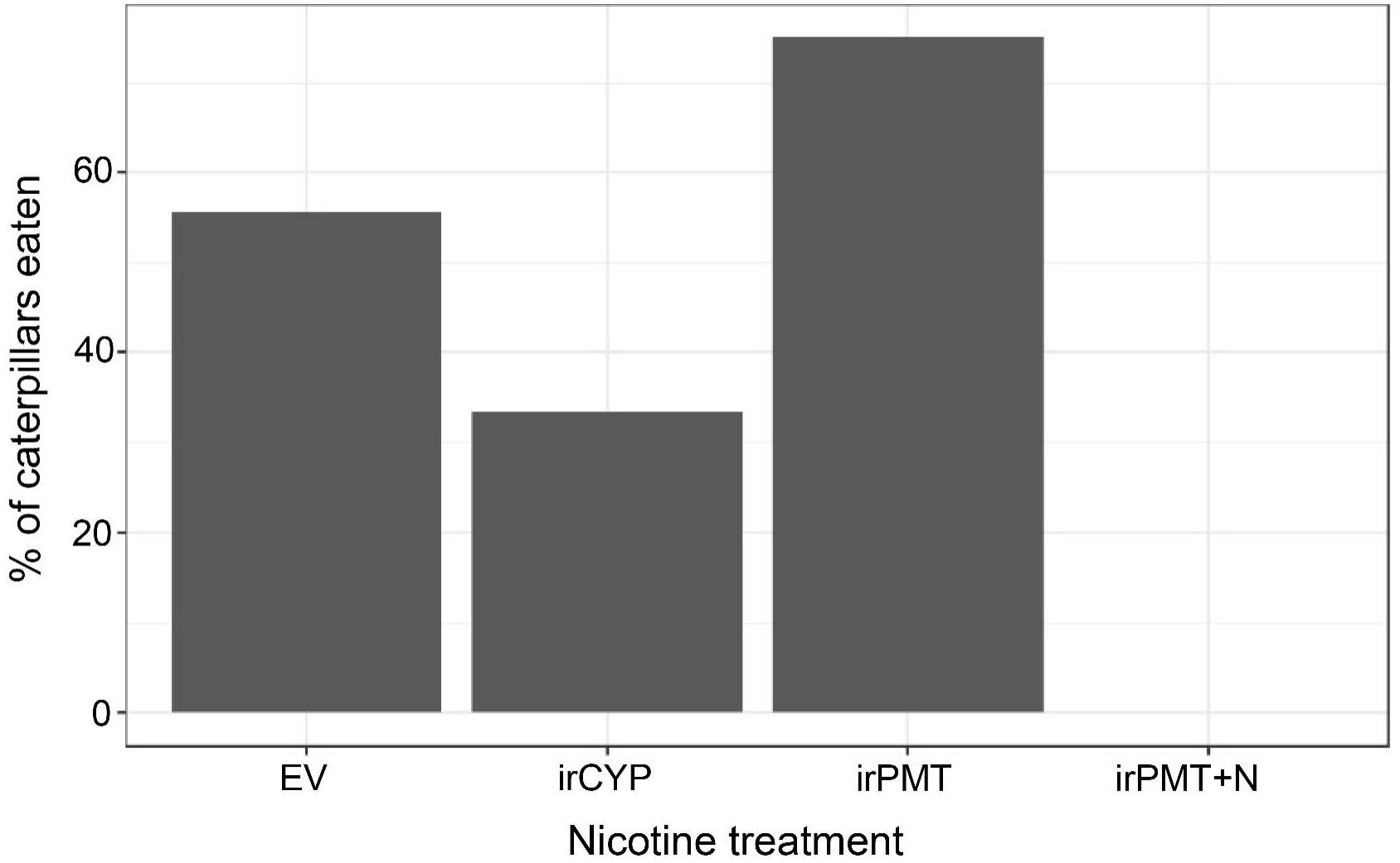
Proportion of *Manduca sexta* trials that resulted in prey being eaten. *Spodoptera* trials are not shown due to 100% of caterpillars being eaten. Full data for both prey species are shown in Table 1.

### Nicotine’s effect on lizard behavior

Most behaviors yielded no significant differences between treatments according to omnibus tests across all treatments (Table 2). Only the rate of two behaviors, lip lick and biting, were found to differ significantly between treatments (Table 3; Figure 3A, 3B). Rates of lip licking were highest during EV and irPMT trials, whereas the PMT+N treatment was associated with the lowest rates of this behavior (Figure 3A; Table 3). With respect to biting, the only significant difference was between the irPMT and irPMT+N treatments, with lower rates of biting directed at the nicotine-sprayed prey. EV and irCYP treatments yielded intermediate bite rates that were not significantly different from any other treatment groups (Figure 3B; Table 3). Although the omnibus test (Kruskal-Wallis) wasn’t statistically significant, we also found a striking trend towards a greater proportion of time spent performing aggressive thrashing behavior during trials with irPMT caterpillars, relative to all other treatments (Table 2; Figure 3C)

**Table 2.** Results from statistical analyses of behavior occurrences during trials. *Spodoptera* trial results are not shown as all yielded non-significant results.

| PREY SPECIES | BEHAVIOR | TEST | STATISTIC<sup>#</sup> | P-VALUE |
|---|---|---|---|---|
| *M. SEXTA* | Bite Rate | ANOVA | *3.136* | *0.039\** |
| *M. SEXTA* | Thrashing Duration | Kruskal-Wallis | 3.060 | 0.383 |
| *M. SEXTA* | Tongue Flick | ANOVA | 1.683 | 0.190 |
| *M. SEXTA* | Lip Lick Rate | Kruskal-Wallis | *11.934* | *0.008\*\** |
| *M. SEXTA* | Head Bob Rate | ANOVA | 1.677 | 0.191 |

<sup>#</sup> Chi-squared for Kruskal-Wallis. F-value for ANOVA.

**Table 3.** *Post-hoc* test results (p-values adjusted for multiple testing) examining pairwise differences between treatments for bite and lip lick occurrence rates during *Manduca sexta* presentation trials.

| **TREATMENT<br>COMPARISON** | **BITES<br>(TUKEY'S HSD)** | **LIP LICKS<br>(CONOVER TEST)** |
|---|---|---|
| *CYP-EV* | 0.963 | 0.064 |
| *CYP-PMT* | 0.135 | *0.038\** |
| *CYP-PMT+N* | 0.862 | 0.855 |
| *EV-PMT* | 0.074 | 0.855 |
| *EV-PMT+N* | 0.990 | *0.029\** |
| *PMT-PMT+N* | *0.045\** | *0.016\** |

**Figure 3.**
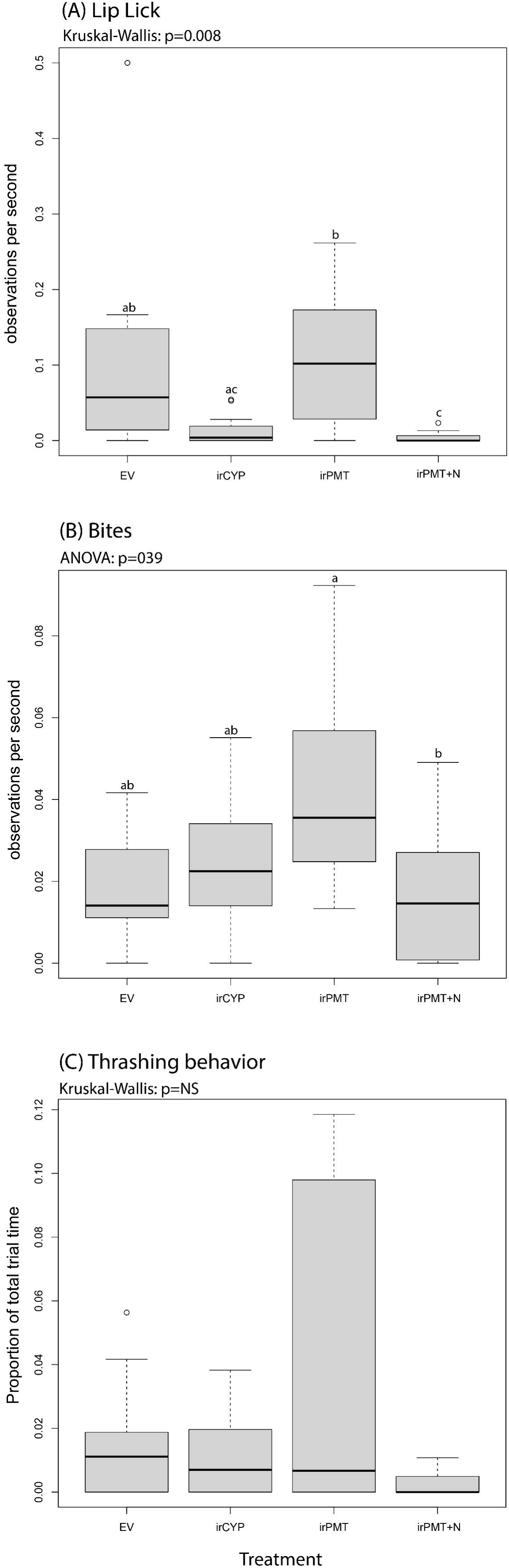
Results of behavior occurrence rates across nicotine treatments. **(A)** shows the rate (observations per second) of lip licking, **(B)** shows the rate of biting pre, and **(C)** shows the proportion of trial time spent thrashing prey.

## Discussion

Chemosensory behaviors, aggressive behaviors, and the likelihood of consuming *M. sexta* larvae were reduced in the presence of nicotine (Figures 2 and 3; Tables 1-3); however, we did not find any evidence to support the hypothesis that defensive halitosis (ie: nicotine exhaled through caterpillar spiracles) prevents predation by lizards (Figure 2; Table 1). In fact, none of our statistical analysis found any differences in lizard behavior towards irCYP and EV (control) caterpillars. However, the increase in chemosensory behaviors directed at nicotine-free irPMT caterpillars (Figure 2A) suggests that lizards are capable of perceiving nicotine. As the trials only began when lizards physically interacted with a caterpillar, they may have perceived surface nicotine present on caterpillar cuticles. The role of surface nicotine for influencing lizard behavior is consistent with the significant reduction in chemosensory behavior, aggression, and consumption of irPMT-fed *M. sexta* that were sprayed with nicotine. Although the small sample sizes thwarted the ability to detect significant differences in consumption rates, the fact that no irPMT+nicotine *M. sexta* were consumed by lizards is a striking result (Figure 2; Table 1) consistent with the inference that nicotine is indeed an aversive chemical stimulus for predatory lizards. Future research with larger sample sizes in more controlled (ie: not field-based) settings are likely to yield more conclusive results.

Lizard responses to *Spodoptera exigua* larvae showed no differences between nicotine treatments. In fact, all *S. exigua* presented to lizards were consumed regardless of their host-plant genotype. Past studies have found that nicotine metabolites, such as cotinine, lack the predation-deterring properties of nicotine (Kumar et al 2014a); as such, the rapid detoxification of nicotine by generalist herbivores could explain the results; however, we also found that lizards consumed the irPMT+nicotine *S. exigua* larvae. This suggests that lizard responses to nicotine are dose dependent. Given that *S. exigua* larvae are significantly smaller than *M. sexta* larvae, it may be that lizards readily consume small prey regardless of nicotine while aversive chemical stimuli play a more prominent role in decisions surrounding the consumption of larger prey insects. Future studies may benefit from the use of size-matched, rather than our use of age-matched larvae, to better assess the responses of lizards to various prey species.

Lizards are known to vary with respect to their foraging strategies, with some species employing a sit-and-wait/ambush strategy and other, typically highly mobile, species actively foraging for prey. Active foragers rely more heavily on their chemical senses to locate prey (Cooper 1995b) and have even been found to attack cotton swabs when they are treated with prey-derived chemical cues (Goldberg et al 2021); something that is rarely observed in sit-and- wait foraging species (Cooper and Van Wyk 1994; Cooper 2000; Goldberg et al 2019). *Uta stansburiana* is a sit-and-wait foraging species (Best and Gennaro 1984), a trait that is undoubtedly influencing its responses to prey-associated chemical cues. An intriguing direction for future work may be to compare the responses of *U. stansburiana* with an actively foraging lizard species (e.g. members of the genus *Aspidoscelis*; Goldberg et al. 2021) in order to better understand the ways that lizard foraging mode influences their sensitivity to toxin-containing prey.

## Supporting information

Supplemental Table 1 - Raw data

## Acknowledgments

We thank Brigham Young University for use of the Lytle Ranch Preserve field station; Danny Kessler, Tanja Bloss, Anne Adler, Meredith Schuman, and Celia Diezel for help with the field experiments; and the Max Planck Institute for Chemical Ecology glasshouse team.

## Permits and Ethics Statement

This study complies with the 4R (reduce, refine, replace, responsibility) animal research guidelines of the Max Planck Society which does not issue formal permits for research with vertebrate animals. Import and release of transgenic plants into the US was done in accordance with multiple Animal and Plant Health Inspection Service (APHIS) permits: 07-341-101n, 11-350-101r, 12-333-101r, 04-020-07n, 04-344-06n, 04-020-08n, 04-344-07n, 07-341-101n, 11-350-101r, 12-333-101r, 10-004-105m, 11-350-101r, and 12-333-101r.

## Data Availability Statement

The raw data that was used in this study are provided as Supplemental Table 1. The video footage underlying the data is available in Goldberg and Baldwin (2026) on Zenodo: https://zenodo.org/records/24439319

## Author Contributions

Caroline Pete: formal analysis, visualization, writing – original draft preparation (lead), writing – review and editing (supporting). Sarah E Pottinger: data curation, writing – review and editing (supporting). Ian T Baldwin: conceptualization (supporting), funding acquisition, methodology (supporting), resources, writing – review and editing (supporting). Jay K Goldberg: conceptualization (lead), methodology (lead), investigation, writing – original draft preparation (supporting), writing – review and editing (lead).

---

**Supplemental Table 1**. Raw data used in statistical analyses

## References

Benowitz, N.L., Hukkanen, J., Jacob, P., 2009. Nicotine chemistry, metabolism, kinetics and biomarkers. Handb Exp Pharmacol 29–60. 10.1007/978-3-540-69248-5_2

Beran, F., Petschenka, G., 2022. Sequestration of plant defense compounds by insects: From Mechanisms to Insect–Plant Coevolution. Annual Review of Entomology 67, 163–180. 10.1146/annurev-ento-062821-062319

Best, T.L., Gennaro, A.L., 1984. Feeding Ecology of the Lizard, Uta stansburiana, in Southeastern New Mexico. Journal of Herpetology 18, 291–301. 10.2307/1564083

Bui, T.N.T., Himanen, S.J., Holopainen, J.K., 2021. Environmentally acquired chemical camouflage affects Pieris brassicae L. host plant selection and orientation behaviour of a larval parasitoid. Arthropod-Plant Interactions 15, 299–312. 10.1007/s11829-021-09830-8

Chang, L., Xu, Z., Deng, P., Zhang, N., He, T., Liu, X., He, W., Zheng, A., Hu, W., Pan, M., Li, W., Halitschke, R., Li, R., Fan, M., Baldwin, I.T., Zhang, Y., Li, D., 2026. Complete biosynthesis of nicotine. Cell 189, 2700–2713.e20. 10.1016/j.cell.2026.03.034

Cooper Jr, W.E., 2000. An adaptive difference in the relationship between foraging mode and responses to prey chemicals in two congeneric Scincid lizards. Ethology 106, 193–206. 10.1046/j.1439-0310.2000.00523.x

Cooper Jr., W.E., van Wyk, J.H., 1994. Absence of prey chemical discrimination by tongue-flicking in an ambush-foraging lizard having actively foraging ancestors. Ethology 97, 317–328. 10.1111/j.1439-0310.1994.tb01050.x

Cooper, W.E., 1997. Correlated evolution of prey chemical discrimination with foraging, lingual morphology and vomeronasal chemoreceptor abundance in lizards. Behav Ecol Sociobiol 41, 257–265. 10.1007/s002650050387

Cooper, W.E., 1995a. Prey chemical discrimination and foraging mode in Gekkonoid lizards. Herpetological Monographs 9, 120–129. 10.2307/1467000

Cooper, W.E., 1995b. Foraging mode, prey chemical discrimination, and phylogeny in lizards. Animal Behaviour 50, 973–985. 10.1016/0003-3472(95)80098-0

Cooper, W.E., 1994. Chemical discrimination by tongue-flicking in lizards: A review with hypotheses on its origin and its ecological and phylogenetic relationships. J Chem Ecol 20, 439–487. 10.1007/BF02064449

Goldberg, J.K., Baldwin, I.T., 2026. Video footage associated with “Nicotine stimulates chemosensory behavior and reduces aggression towards tobacco-feeding caterpillars (Manduca sexta) by predatory side-blotched lizards (Uta stansburiana)” Zenodo 10.5281/zenodo.24439319

Goldberg, J.K., Pintel, G., Pruett, J.A., Weiss, S.L., 2021. Whiptail lizards (Aspidoscelis exsanguis) recognize invertebrate prey via cuticular hydrocarbons. Ethology 127, 416–423. 10.1111/eth.13142

Goldberg, J.K., Pintel, G., Weiss, S.L., Martins, E.P., 2019. Predatory lizards perceive plant-derived volatile odorants. Ecology and Evolution 9, 4733–4738. 10.1002/ece3.5076

Kessler, D., Bhattacharya, S., Diezel, C., Rothe, E., Gase, K., Schöttner, M., Baldwin, I.T., 2012. Unpredictability of nectar nicotine promotes outcrossing by hummingbirds in Nicotiana attenuata. Plant J 71, 529–538. 10.1111/j.1365-313X.2012.05008.x

Kumar, P., Pandit, S.S., Steppuhn, A., Baldwin, I.T., 2014a. Natural history-driven, plant-mediated RNAi-based study reveals CYP6B46’s role in a nicotine-mediated antipredator herbivore defense. Proceedings of the National Academy of Sciences 111, 1245–1252. 10.1073/pnas.1314848111

Kumar, P., Rathi, P., Schöttner, M., Baldwin, I.T., Pandit, S., 2014b. Differences in nicotine metabolism of two Nicotiana attenuata herbivores render them differentially susceptible to a common native predator. PLOS ONE 9, e95982. 10.1371/journal.pone.0095982

Martins, E.P., Labra, A., Halloy, M., Tolman Thompson, J., 2004. Large-scale patterns of signal evolution: an interspecific study of Liolaemus lizard headbob displays. Animal Behaviour 68, 453–463. 10.1016/j.anbehav.2003.08.026

Narberhaus, I., Zintgraf, V., Dobler, S., 2005. Pyrrolizidine alkaloids on three trophic levels – evidence for toxic and deterrent effects on phytophages and predators. Chemoecology 15, 121–125. 10.1007/s00049-005-0302-z

Pomerleau, O.F., 1992. Nicotine and the central nervous system: Biobehavioral effects of cigarette smoking. The American Journal of Medicine, The Effects of Cigarette Smoking: A Global Perspective 93, S2–S7. 10.1016/0002-9343(92)90619-M

R Core Team 2025. R: A Language and Environment for Statistical Computing. R Foundation for Statistical Computing, Vienna, Austria. https://www.R-project.org/

Santis, G.D., Okura, Y., Hirata, K., Ishiuchi, S.-I., Fujii, M., Xantheas, S.S., 2024. Affinity of nicotinoids to a model nicotinic acetylcholine receptor (nAChR) binding pocket in the human brain. J. Phys. Chem. B 128, 4577–4589. 10.1021/acs.jpcb.3c07919

Steppuhn, A., Gase, K., Krock, B., Halitschke, R., Baldwin, I.T., 2004. Nicotine’s defensive function in nature. PLOS Biology 2, e217. 10.1371/journal.pbio.0020217

Stork, W.F.J., Weinhold, A., Baldwin, I.T., 2011. Trichomes as dangerous lollipops: Do lizards also use caterpillar body and frass odor to optimize their foraging? Plant Signaling & Behavior 6, 1893–1896. 10.4161/psb.6.12.18028

Xu, S., Brockmöller, T., Navarro-Quezada, A., Kuhl, H., Gase, K., Ling, Z., Zhou, W., Kreitzer, C., Stanke, M., Tang, H., Lyons, E., Pandey, P., Pandey, S.P., Timmermann, B., Gaquerel, E., Baldwin, I.T., 2017. Wild tobacco genomes reveal the evolution of nicotine biosynthesis. Proceedings of the National Academy of Sciences 114, 6133–6138. 10.1073/pnas.1700073114

